# Evaluation of the SARS-CoV-2 inactivation efficacy associated with buffers from three kits used on high-throughput RNA extraction platforms

**DOI:** 10.1101/2021.04.14.439928

**Authors:** Ruth E. Thom, Lin S. Eastaugh, Lyn M. O’Brien, David O. Ulaeto, James S. Findlay, Sophie J. Smither, Amanda L. Phelps, Helen L. Stapleton, Karleigh A. Hamblin, Simon A. Weller

**Affiliations:** CBR Division, Dstl Porton Down, Salisbury, United Kingdom

## Abstract

Rapid and demonstrable inactivation of SARS-CoV-2 is crucial to ensure operator safety during high-throughput testing of clinical samples. The inactivation efficacy of SARS-CoV-2 was evaluated using commercially available lysis buffers from three viral RNA extraction kits used on two high-throughput (96-well) RNA extraction platforms (Qiagen QiaCube HT and the ThermoFisher Kingfisher Flex) in combination with thermal treatment. Buffer volumes and sample ratios were chosen for their optimised suitability for RNA extraction rather than inactivation efficacy and tested against a representative sample type; SARS-CoV-2 spiked into viral transport medium (VTM). A lysis buffer from the MagMax Pathogen RNA/DNA kit (ThermoFisher), used on the Kingfisher Flex, which included guanidinium isothiocycnate (GITC), a detergent, and isopropanol demonstrated a minimum inactivation efficacy of 1 x 10^5^ TCID_50_/ml. An alternative lysis buffer from the MagMax Viral/Pathogen Nucleic Acid kit (Thermofisher) also used on the Kingfisher Flex and the lysis buffer from QIAamp 96 Virus QIAcube HT Kit (Qiagen) used on the QiaCube HT (both of which contained GITC and a detergent) reduced titres by 1 x 10^4^ TCID_50_/ml but did not completely inactivate the virus. Heat treatment alone (15 minutes, 68 °C) did not completely inactivate the virus, demonstrating a reduction of 1 x 10^3^ TCID_50_/ml. When inactivation methods included both heat treatment and addition of lysis buffer, all methods were shown to completely inactivate SARS-CoV-2 inactivation against the viral titres tested. Results are discussed in the context of the operation of a high-throughput diagnostic laboratory.

## INTRODUCTION

Severe acute respiratory syndrome coronavirus-2 (SARS-CoV-2) belongs to the Coronaviridae family and is the causative agent of the respiratory illness, coronavirus disease (COVID-19) (1). The enveloped positive-sense single-stranded RNA virus was first discovered in early 2020 after a cluster of viral pneumonia cases of unknown cause were reported in the Hubei Province of China (2). The virus is highly contagious in humans and in March 2020 The World Health Organisation (WHO) declared a global pandemic (3).

Diagnostic testing is critical in the fight against the COVID-19 pandemic (4), not just for patients displaying symptoms but also for asymptomatic carriers and pre-symptomatic patients (5). SARS-CoV-2 has been classified in the UK as a Hazard Group (HG) 3 pathogen by the Advisory Committee for Dangerous Pathogens (ACDP), meaning that this virus must be handled under Containment Level (CL) 3 conditions. However, guidance from WHO (6) and Public Health England, UK (7) has permitted non-propagative diagnostic testing to be carried out at CL 2 with non-inactivated samples being handled within a Class I microbiology safety cabinet.

Real-time reverse transcriptase polymerase chain reaction (RT-PCR) is the gold standard test to for the detection of SARS-CoV-2 from nasopharyngeal swab samples (8). Inactivation of viral pathogens prior to PCR is typically carried out at the same time as extraction of viral nucleic acids from samples, with chemical or physical methods employed. Typically buffers provided in nucleic acid extraction kits contain chaotropic salts, solvents, and detergents to lyse the virus. Guanidinium salts, such as guanidinium thiocyanate (GITC), are chaotropic agents found in many lysis buffers which in some cases have been demonstrated to inactivate viral pathogens, including alphaviruses, flaviviruses, filoviruses and a bunyavirus (9, 10). Other reports though suggest that a combination of a GITC containing extraction buffer (such as Qiagen AVL) and a solvent (such as ethanol), is required for the inactivation of viruses such as Ebola virus (11) and Middle East Respiratory Syndrome coronavirus (MERS-CoV) (12). Detergents such as Tween, SDS and Triton X100 have also been shown to disrupt viral envelopes and reduce viral titres (13–15), with a combination of the GITC based reagent (Buffer AVL) and Triton X100 having been reported to inactivate Ebola virus (16). Physical processes such as heat can also be incorporated in the nucleic acid extraction workflow and can have an inactivation effect. Some reports suggest that the application of heat alone can inactivate SARS-CoV, MERS-CoV and SARS-CoV-2 following a heat regimen of 65 °C for at least 15 minutes (17–19).

Due to commercial sensitivity, manufacturers of extraction kits are not required to publish the full ingredient list of proprietary buffers (with potential viral inactivating components only inferred if they are listed on associated Material Safety Data Sheets(MSDS)) and post-treatment viability test methods vary in stringency across studies. Due to the disparate and varying literature sources describing the efficacy of inactivation methods for the extraction of RNA, a standardised protocol for the inactivation of SARS-CoV-2 was developed and experimental validation of the different approaches was undertaken.

Since the pandemic was declared, UK’s Defence Science Technology Laboratory (Dstl) and British military clinicians have set up the Defence COVID lab (DCL), which has been awarded an extension to scope (under ISO17025) for the provision of a SARS-COV-2 PCR test by the United Kingdom Accreditation Service (UKAS). The DCL analyses samples from UK military units and operates two automated high-throughput RNA extraction platforms (Qiagen QiaCube HT and the ThermoFisher Kingfisher Flex). In this study we report the inactivation efficacy of SARS-CoV-2 by buffers from three commercially available kits used on these two platforms. Buffer volumes and ratios were chosen for their suitability for RNA extraction (following manufacturer’s instructions) rather than their potential inactivation efficacy, however in doing so we have further investigated the inactivation efficacy of combinations of GITC containing buffers, solvents, and/or detergents with and without an additional heat inactivation step. We provide evidence to support protocols for the inactivation of SARS-CoV-2 and the safe use of clinical samples in down-stream RT-PCR in high-throughput diagnostic laboratories.

## METHODS

### Virus strains, cell culture and reagents

All cell culture was carried out using confluent monolayers of Vero C1008 cells (European Collection of Cell Cultures [ECACC], United Kingdom; catalogue no. 85020206) maintained in Dulbecco’s minimal essential medium (DMEM; Sigma, United Kingdom) supplemented with 10% fetal calf serum, 1% L-glutamine and 1% penicillin-streptomycin (Sigma, United Kingdom) and incubated at 37 °C in a 5% CO_2_ environment. Prior to virus being added to cell monolayers, 10% DMEM was replaced with Leibovitz’s L-15 (to buffer for the lack of CO_2_ at CL3), supplemented as described for DMEM, with the exception of 2% fetal calf serum and incubated at 37 °C. All virus manipulations were carried out under BSL/CL 3 conditions using the SARS-CoV-2 England 2 strain (GISAID reference EPI_ISL_407073), provided by Public Health England. Virus stock was propagated in Vero C1008 cell, harvested at day 3 and clarified by centrifugation at 350 x g for 15 minutes (Sigma 3-16K centrifuge). Viral stocks were concentrated by centrifugation at 11, 000 x g for 3 hours at 4 °C to achieve 1×10^8^ Tissue Culture Infectious Dose (TCID) _50_/ml. All virus stocks were stored at −80 °C. Buffers and reagents from three different RNA extraction kits were assessed to determine inactivation of SARS-CoV-2 (Table 1). The composition of these initial reagents and their suitability for extraction of SARS-CoV-2 RNA from clinical samples was determined based on manufacture protocols and after discussions with each manufacturer.

**TABLE 1.** Protocols tested for assessing inactivation using lysis buffers.

| Manufacturer,<br>RNA extraction kit,<br>Platform. | Reagents<br>(volume / sample) | Active virucidal<br>components* | Reagent :<br>Sample ratio |
| --- | --- | --- | --- |
| Qiagen,<br>QIAamp 96 Virus QIAcube<br>HT Kit,<br>Qiagen Qiacube HT.<br>(Referred to here as Qiagen<br>protocol) | ACL buffer (190 µl) | GITC 30 - <50% | 1.6 : 1 |
|  | ATL buffer (100 µl) | 1 - <3% SDS |  |
|  | Proteinase K (20 µl) |  |  |
|  | Carrier RNA (5 µl) |  |  |
|  | MS2 (10 µl) |  |  |
| ThermoFisher,<br>MagMax Pathogen<br>RNA/DNA kit,<br>Kingfisher Flex. | Lysis binding<br>buffer (350 µl) | GITC 55-80%<br><0.001%<br>Acrylamide<br>Zwittergent | 3.8 : 1 |
| *(Referred to here as*<br>*MagMax Protocol 1)* | Isopropanol (300<br>µl) | 100% 2-propanol |  |
|  | Carrier RNA (2 µl) |  |  |
|  | Water (100 µl) |  |  |
|  | MS2 (10 µl) |  |  |
| ThermoFisher,<br>MagMax viral/pathogen<br>nucleic acid isolation kit,<br>Kingfisher Flex.<br>*(Referred to here as*<br>*MagMax Protocol 2)* | Lysis binding<br>buffer (265 µl) | GITC 55-80%<br><0.001%<br>Acrylamide<br>Zwittergent | 1.4 : 1 |
|  | Proteinase K (5 µl) |  |  |
|  | †Water (Magnetic<br>beads) (10 µl) |  |  |
|  | MS2 (10 µl) |  |  |

\*As identified directly from components, manufacturer information, or inferred from the associated MSDS.
†Water was used to replace the magnetic beads as the washing steps described below would not remove the beads and the beads interfered the read-out of the TCID<sub>50</sub> assay.
GITC: Guanidinium thiocyanate. SDS: Sodium dodecyl sulphate

### Viral inactivation

The inactivation efficacy of the lysis buffers in all three protocols was evaluated with and without the inclusion of a heat step. Table 1 summarises the components and volumes used for each lysis buffer preparation. MS2 bacteriophage (10^6^ Plaque Forming Unit (PFU)/ml) was added to each lysis buffer preparation as an internal control in the DCL, Dstl. Test samples for each experiment were set up in triplicate and each experiment was performed on at least three separate occasions.

Viral transfer medium (VTM; EO Labs, United Kingdom) was inoculated with SARS-CoV-2 to achieve a starting concentration of 5×10^6^ TCID_50_/ml for all experiments. To the lysis buffer preparations, 200 μl of virus in VTM was added, the samples were briefly vortexed and incubated for 10 minutes at room temperature. For heat treated samples, the tubes were incubated for 25 minutes in a heat block (Eppendorf ThermoMixer C heat) set at 75 °C. Laboratory tests showed that this was the temperature setting required for this individual heat block to heat and maintain the samples at 68 °C for 15 minutes. Heat steps were carried out after the addition of virus to either lysis buffer reagents or to an equivalent volume of tissue culture medium (TCM), to assess the effect of viability following heat in the presence or absence of reagents. Further controls included sham-inactivated virus, where appropriate volume of TCM replaced the lysis buffer reagents and negative controls consisting of VTM-only added to lysis buffer reagents to assess the effect of the reagents on cell monolayers.

After inactivation (with or without heat treatment) all samples and controls were pelleted by centrifugation at 6, 000 x *g* for 5 minutes in a microcentrifuge (Hermle Microlitre Centrifuge Z 160 M). The supernatant was discarded and the pellet resuspended in 1 ml TCM and washed a further 4 times for the Qiagen reagents and 2 times for the Kingfisher reagents in order to remove all traces of the inactivation chemicals from the sample and to avoid toxicity during cell culture. After the final wash the pellets were re-suspended in 1ml of TCM.

### Post inactivation viral viability assays

To quantify and determine the viability of the virus following inactivation, the samples were prepared for TCID_50_ end-point dilution assay (20) and the remaining sample underwent three rounds of serial passage in tissue culture flasks.

In brief, TCID_50_ assay was performed using Vero C1008 cells prepared in 96-well microtitre plates to achieve confluent monolayers on the day of assay. To all wells of column 1 of the plate 100 μl of test sample was added. From column 1, 20 μl of sample was transferred sequentially across the plate to achieve a 10-fold serial dilution to column 9. Cells in columns 11 and 12 were left in TCM as controls. Plates were incubated in a humidified atmosphere for 3 - 4 days at 37 °C, after which they were scored for cytopathic effects (CPE) by microscopic observation. The TCID_50_ value was calculated by the method of Reed and Muench (21).

For secondary confirmation of viral inactivation, all of the remaining sample (approx. 180 μl) was added to confluent monolayer of Vero C1008 cells in a 12.5 cm^2^ tissue culture flask. Flasks were incubated in a humidified atmosphere for 3 - 4 days after which presence or absence of cytopathic effect was recorded. A total of three passages were performed and CPE recorded after each round. To control for cross-contamination a set of un-infected flasks were also prepared and supernatant passaged in parallel to the experimental samples. A 10-fold serially dilution of SARS-CoV-2 was also inoculated into a set of flasks starting from 1.7 x 10^7^ TCID_50_/ml and diluted to 1.1 TCID_50_/ml to show the Limit of Detection (LOD) of the flask passage assay and demonstrate a suitable environment for the passage and propagation of the virus.

### Statistical analysis

All data were graphically represented and statistically analysed using GraphPad Prism 8. Kruskal-Wallis analysis of variance (ANOVA) was performed on data sets with Dunn’s multiple comparison post hoc.

## RESULTS

The inactivation of SARS-CoV-2 was assessed using three different RNA lysis buffers with and without the inclusion of a heat step. The viability of virus was determined quantitatively using the TCID_50_ assay and qualitatively by serially passaging samples in flask.

### Determination of starting concentration of SARS-CoV-2

These studies used the highest working concentration of SARS-CoV-2 that was available and this ranged from 5.9×10^5^ to 3.5×10^6^ TCID_50_/ml (Figure 1). Following the inactivation procedure residual toxic lysis buffer components were removed by way of multiple wash steps. Residual chemical components would otherwise be toxic to the cell based assays. To determine if the multiple wash steps by centrifugation resulted in a loss of virus, virus was inoculated into TCM without the addition of lysis reagents (as described in materials and methods) and assayed as described. This highlighted there was approximately a 1-Log_10_ drop in titre, providing a mean viral titre of 2.4×10^5^ TCID_50_/ml (Figure 1A, B and C).

**FIGURE 1.**
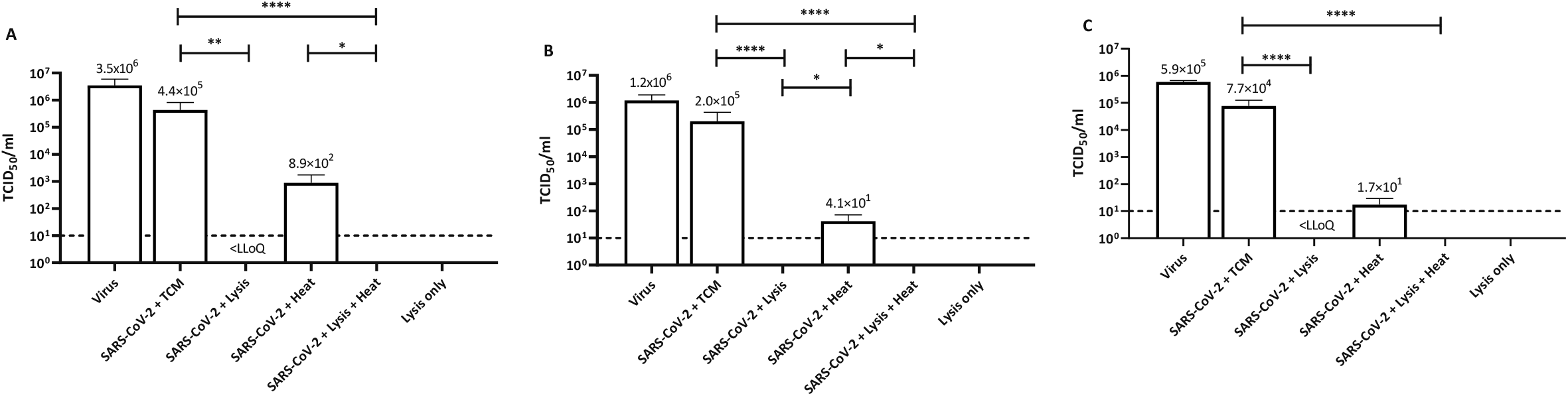
Titre of SARS-CoV-2 by TCID_50_ assay following inactivation protocols. A. Qiagen protocol, B. MagMax protocol 1, B. MagMax protocol 2. Mean + Standard Deviation collated from triplicate results from three separate occasions (n=9). Dashed line = Lower limit of quantification (LLoQ < 10 TCID_50_/ml); Tissue culture media (TCM). Kruskal-Wallis ANOVA with Dunn’s multiple comparison post hoc, where * p <0.05, **p<0.01, ***p<0.001, ****p<0.0001; statistical analysis excludes virus stock and lysis only data.

### Chemical inactivation of SARS-CoV-2

When virus was added to the Qiagen lysis buffer there was a statistically significant 5-Log_10_ drop in virus titre from 4.4×10^5^ TCID_50_/ml to below the lower limit of quantification (LLoQ) (p=0.002) Complete inactivation was not achieved however, as virus was detected below the LLoQ but this was not quantifiable. However by extrapolation it was estimated that the titre was 6.2 TCID_50_/ml (Figure 1A).

Similar results were observed when virus was inactivated using the MagMax protocol 2; complete inactivation was not achieved as virus was detected below the LLoQ, and was not quantifiable. The starting titre of virus for these experiments, following washing steps was 7.7×10^4^ TCID_50_/ml, demonstrating a 4-Log_10_ drop in viral titre following inactivation (p<0.0001) (Figure 1C).

Virus inactivation following the MagMax protocol 1 resulted in no detectable virus by TCID_50_ assay. The starting concentration of virus, following washing steps was calculated to be 2×10^5^ TCID_50_/ml, thus demonstrating a 5-Log_10_ drop in viral titre with this particular protocol (p<0.0001) (Figure 1B).

### Heat inactivation of SARS-CoV-2

Heat alone or in combination with lysis buffer was also investigated as a means to inactivate SARS-CoV-2. For each experiment, virus in TCM was heated at 68 °C for 15 minutes and centrifuged to maintain consistency with samples in lysis buffer. Although not statistically significant, at least a 3-Log_10_ drop in viral titre was observed following heat treatment alone, with an average titre of remaining viable virus across all experiments of 3.2×10^2^ TCID_50_/ml (Figure 1).

When the virus was added to either of the three lysis buffers and subsequently heated, no viable virus was detected following TCID_50_ assay and an average drop in viral titre of 5-Log_10_ across all experiments (p<0.0001) (Figure 1A, B and C).

### Confirmation of inactivation by viral propagation

To confirm findings by TCID_50_ assay viral samples were propagated in cell culture flasks over a total of three passages to identify potential viral break-through. Table 2 shows the results of the presence of CPE after the first passage. The limit of detection for viral propagation was determined following propagation of serially diluted virus stocks (Table 2 row 1 to 5) and on average the limit of detection was 1.3 TCID_50_/ml.

**TABLE 2.** Summary of results following cell culture passage and TCID_50_ assay. Passage results shown are after the third serial. TCID_50_ titres are mean titre/ml and standard deviation. * Indicates the TCID_50_/ml is extrapolated from known starting concentration and calculated based on number of flasks infected. SARS-2 = SARS-CoV-2. TCM = Tissue culture media. LLoQ = Lower limit of Quantification (< 10 TCID_50_/ml). SD = Standard deviation.

| Inactivation protocol | Qiagen protocol |  | MagMax protocol 1 |  | MagMax protocol 2 |  |
| --- | --- | --- | --- | --- | --- | --- |
| SAMPLE DESCRIPTION | Flasks | TCID<sub>50</sub>/ml | Flasks | TCID<sub>50</sub>/ml | Flasks | TCID<sub>50</sub>/ml |
|  | infected/<br>total flasks | (SD) | infected/<br>total flasks | (SD) | infected/<br>total flasks | (SD) |
| 1. SARS-CoV-2 Starting titre | 3/3 | 1.7 x 10<sup>7</sup><br>(1.2 x 10<sup>7</sup>) | 3/3 | 5.9 x 10<sup>6</sup><br>(3.6 x 10<sup>6</sup>) | 3/3 | 3.0 x 10<sup>6</sup><br>(3.8 x 10<sup>5</sup>) |
| 2. SARS-CoV-2 10<sup>-4</sup> dilution | 3/3 | 1.7 x 10<sup>3</sup> * | 3/3 | 5.9 x 10<sup>2</sup> * | 3/3 | 3.0 x 10<sup>2</sup> * |
| 3. SARS-CoV-2 10<sup>-5</sup> dilution | 3/3 | 1.7 x 10<sup>2</sup> * | 3/3 | 59.4 * | 2/3 | 20.0 * |
| 4. SARS-CoV-2 10<sup>-6</sup> dilution | 3/3 | 17 * | 1/3 | 2.0 * | 1/3 | 0.7 * |
| 5. SARS-CoV-2 10<sup>-7</sup> dilution | 2/3 | 1.1 * | 0/3 | 0 * | 0/3 | 0 * |
| 6. SARS-CoV-2 + TCM | 9/9 | $4.4 \times 10^5$<br>($3.8 \times 10^5$) | 9/9 | $2.0 \times 10^5$<br>($2.3 \times 10^5$) | 9/9 | $7.7 \times 10^4$<br>($4.8 \times 10^4$) |
| 7. SARS-CoV-2 + lysis buffer | 3/9 | <LLOQ | 0/9 | 0 | 0/9 | <LLOQ |
| 8. SARS-CoV-2 + heat | 9/9 | $8.9 \times 10^2$<br>($8.5 \times 10^2$) | 9/9 | 41.4<br>(30.0) | 8/9 | 17.4<br>(12.1) |
| 9. SARS-CoV-2 + lysis buffer +<br>heat | 0/9 | 0 | 0/9 | 0 | 0/9 | 0 |
| 10. TCM + Lysis buffer | 0/9 | 0 | 0/0 | 0 | 0/9 | 0 |

When virus was added to TCM, CPE was present in all flasks as expected (Table 2 row 6, positive control). No cell toxicity was observed from negative control samples were TCM-only was added to lysis buffer and washed as described previously (Table 2 row 10, negative control).

When SARS-CoV-2 was inactivated following the Qiagen protocol, 3 out of the 9 flasks were scored as positive for CPE. Of the flasks where no CPE was observed, no break-through of virus was seen as a result of serial passage (Table 2 row 7). This data aligns with the TCID_50_ assays, where Qiagen lysis buffer alone did not completely inactivate the virus. Following both MagMax protocols, 0 out of the 9 flasks were scored positively for CPE (Table 2 row 7). For the MagMax protocol 1 this confirms the TCID_50_ results, where no viable virus was also observed. For the MagMax protocol 2, virus was detected but not quantifiable in the TCID_50_ assay (below the LLoQ), however subsequent serial passage did not provide evidence of viability, as all flasks were negative for CPE.

When SARS-CoV-2 was added to TCM and heated for 15 minutes at 68 °C, CPE was observed in all but one flask (Table 2 row 8) confirming the TCID_50_ results that the heating protocol described here does not completely inactivate the virus.

For all inactivation protocols, when SARS-CoV-2 samples were treated in a two-step manner, (lysis buffer and heat), no viable virus was detected in either the quantitative or qualitative assays (Figure 1 and Table 2 row 9). This data provides strong evidence that the lysis buffers described here in combination with the heat protocol can completely inactivate up to 5-Log_10_ TCID_50_/ml SARS-CoV-2.

## DISCUSSION

Real-time PCR is the gold standard clinical diagnostic method for the detection of SARS-CoV-2 in patients displaying symptoms of COVID-19. There has been a rapid development in RNA extraction and RT-PCR diagnostic methods in order to help prevent further spread of infection through communities. It is crucial that testing is accurate and efficient, both of which must not compromise safety of those processing the samples (22). Laboratory acquired infections due to incomplete inactivation or incorrect handling of samples have been reported for SARS-CoV (23, 24) as well as many other infectious agents (25). To date, there are only a handful of publications reporting the use of nucleic acid isolation reagents, detergents and heat to inactivate SARS-CoV-2 (18, 26-28).

In our study we investigated the SARS-CoV-2 inactivation efficacy of viral lysis buffers from three commercially available kits developed to allow RNA extraction on high-throughput (96 well) automated platforms. For each kit the initial lysis buffer mix, developed from manufacturer’s instructions, included a guanidine based lysis buffer with additional viral inactivating components such as a solvent and / or a detergent. Each mix was added to 200 μl of a representative clinical sample (SARS-CoV-2 in viral transport medium). Furthermore we tested all three protocols with and without the addition of a thermal inactivation step at 68 °C for 15 minutes.

We started with the highest possible titre of SARS-CoV-2 that we had available and first determined the titre of virus following wash steps, which were required to remove any chemical compounds that would be cytotoxic to the cell based assays. We chose to remove the reagents from the samples by centrifugation and in doing so, demonstrated a loss of approximately 1-Log_10_ of virus. Other researchers have used centrifugation columns or filters but again report a similar loss in viral titre (14) or residual toxicity leading to reduced sensitivity of the read-out of the assays (28). The wash steps employed here eliminated all residual toxicity, allowing the sensitivity of our assay read-outs to be unaffected.

In our study, the chemicals used to assess the inactivation of SARS-CoV-2 were combinations of GITC, detergent and solvent. The Qiagen protocol (using reagents from the QIAamp 96 Virus QIAcube HT Kit) and the MagMax Protocol 2 (using reagents from the MagMax viral/pathogen nucleic acid isolation kit) both included GITC and a detergent, (SDS or zwittergent, respectively) (Table 1). Both of these inactivation buffers significantly reduced viral titres of SARS-CoV-2 by 4-Log_10_ however complete inactivation of viable virus was not achieved as detectable, but not quantifiable, virus was detected in the TCID_50_ assay (below LLoQ). Subsequent serial passage of viral samples following inactivation using the Qiagen protocol demonstrated virus break-through confirming the results observed in the TCID_50_ assay. It was also anticipated that serial passage of virus inactivated following MagMax protocol 2 would have amplified and enabled virus break-through too, but this was not observed. The stated GITC composition of Qiagen Buffer ACL (30-50%) is lower than that of the MagMax Lysis buffer (55-80%) and thus the higher GITC composition in the MagMax buffer may have exerted a greater efficacy of viral inactivation, although we could not demonstrate complete inactivation. As described previously GITC based chemicals alone have been reported to inactivate some viruses (9, 10) but as observed here and by others this is not always the case (11, 12, 16). Studies by Pastorino *et al* (27), have assessed the inactivation of SARS-CoV-2 using the detergent containing Buffer ATL and in contrast to our findings reported greater than a 6-Log_10_ drop in virus titre. The SDS composition of Buffer ATL used by Pastorino *et al* (2020) was 1 – 10%, however, the SDS composition of ATL buffer in our study, was 1 - <3% SDS (Table 1). Pastorino *et al* (2020) also used a 1:1 ratio of ATL buffer to sample, where as in our protocol we used a reagent to sample ratio of 0.5: 1. Thus the work of Pastorino *et al* (2020) infers a higher concentration of this detergent and larger reagent to sample ratio would be critical for the inactivation process. This also underlines the potential for different concentrations of components in products that are ostensibly the same. Patterson *et al* (2020) and Welch *et al* (14, 28) screened a number of detergents for their inactivation efficacy against SARS-CoV-2. Patterson *et al* (2020) reported that 0.5% SDS inactivated SARS-CoV-2, but used a low starting titre of 10^2^ PFU (14), whereas Welch *et al* (2020) also reported a drop in virus titre of 6.5 Log_10_ TCID_50_/ml but viable virus was still observed (28).

In our study, the only protocol that inactivated virus without an additional heat step was MagMax Protocol 1 (using reagents from the MagMax Pathogen RNA/DNA kit), where no CPE was observed from either TCID_50_ assay or following three rounds of serial passage in tissue culture flasks. The MagMax Protocol 1 included the MagMax lysis binding buffer which contained GITC and the detergent Zwittergent. With the addition of 2-propanol within the lysis buffer mix there were, therefore, three components likely to exert a disruptive effect on the SARS-CoV-2 viral envelope. The reagent to sample ratio of 3.8: 1 was also higher, with more than double the volume of lysis buffer mix added to each sample, compared to the other two methods assessed (Table 1).

Our results suggest that both a high reagent to sample ratio and the incorporation of a solvent improved the inactivation efficacy of a chemical only method. The SARS-CoV-2 inactivation efficacy of the GITC-based Buffer AVL (Qiagen) in combination with ethanol has been assessed in two studies. Complete SARS-CoV-2 inactivation was reported by Welch *et al* (2020) (28) in contrast to incomplete inactivation by Pastorino *et al* (2020) (27). This contradiction in findings could be due to the ratios of reagent, solvent and sample used. Both studies used 4 volumes of AVL to 1 volume of sample; however volumes of ethanol used in combination with Buffer AVL may explain the varying results. Welch *et al* (2020) used 4 volumes of ethanol in combination with AVL and sample, whereas Pastorino *et al* (2020) only added 1 volume of ethanol to the AVL-sample combination. In our studies using the MagMax Protocol 1 the ratio of lysis buffer and isopropanol were considerably less with 1.8 volumes of lysis buffer and 1.5 volumes of solvent, but the addition of the detergent Zwittergent (within the MagMax Lysis Buffer) may have enhanced the inactivation. The addition of the enzyme Proteinase K in both the Qiagen method and MagMax protocol 2 did not appear to have enhanced inactivation efficacy.

We also investigated the efficacy of thermal inactivation, by heating the sample to, and then maintaining at, 68 °C for 15 minutes. Heat inactivation alone reduced the viral titre by 3-Log_10_, although this was not statistically significant compared to the controls, and was not as effective as the use of lysis buffers alone. Burton *et al* 2021 (26) report similar findings with incomplete inactivation of SARS-CoV-2 at 56 and 60 °C for up to 60 minutes. In contrast, some studies have reported the successful use of heat for complete inactivation of SARS-CoV and SARS-CoV-2 (17, 18). Kim *et al* 2020 (18) demonstrated the complete inactivation of SARS-CoV-2 in clinical samples following incubation at 65 °C for 30 minutes, although this work was based on quantitativeTCID_50_ assays alone. Furthermore, Darnell *et al* (17) reported complete inactivation of SARS-CoV after heating at 65 °C for 60 minutes, the longer time was required to ensure any viral aggregates were fully exposed and inactivated by the heat treatment.

The use of heat to inactivate virus has been reported to reduce viral RNA stability (29, 30) and depending on the target gene used for RT-PCR, incubation at 65 °C for 30 minutes can significantly reduce the target copy numbers leading to false negative results of clinical samples (18, 30). The DCL has an accredited SARS-CoV-2 diagnostic workflow (31) using the Qiagen and Kingfisher (using MagMax protocol 1) extraction platforms each with an additional heat inactivation step. Multiple External Quality Assessment panels and reference standards have been tested during DCL set-up and operation. The E-Gene PCR assay (32) is used in this laboratory and in our hands the heat inactivation regime we employ does not appear to adversely affect PCR results.

In determining the practical relevance of our work the viral loads in COVID19 samples likely to be encountered in a high-throughput diagnostic laboratory should be considered. Currently there is little information on the infectious viral load present on a clinical nasal/throat swab. Most of the data report Ct values following RT-PCR (33) but one study has estimated that there is a median titre of 10^3^ TCID_50_/ml collected from 90 nasopharyngeal or endotracheal clinical samples (34). During DCL validation studies a precisely defined reference standard dilution series of entire SARS-CoV-2 virions (SARS-CoV-2 Analytical Q Panel; Qnostics Ltd, UK) was tested (data not shown). Within this series the highest concentration of material was 6 Log_10_ digital copies (dC)/ml and following RNA extraction using the Qiagen method described in this paper mean E-gene (32) quantification cycle (C_q_) values of 22.65 were returned from this concentration. During DCL operation we have commonly tested positive samples with E-gene PCR C_q_ values in the low teens, with occasional samples returning C_q_ values <13. Although care must be taken in comparing and extrapolating PCR (C_q_), TCID_50_/ml and dC/ml values this is consistent with a study reporting similarly low C_q_ values from COVID patients early in the infection cycle (35) and indicates that some swab samples can contain very high viral loads.

We have demonstrated the SARS-CoV-2 inactivation efficacy of the reagents found in lysis buffers of three commercially available kits used on high-throughput extraction platforms. Only when combined with a heat step did all methods show a complete inactivation of SARS-CoV-2 by both TCID_50_ assay and by sequential passage in tissue culture. Therefore in the DCL samples are sequentially mixed with lysis buffer and then followed with heat treatment. This approach also extends the contact time of lysis buffer to sample which should further enhance the inactivation efficacy of the buffers and mitigates the fact that in this inactivation study we were unable to test samples with a starting concentration greater than 5.9×10^5^ TCID_50_/ml (in view of the likely higher concentrations seen in samples received). In our studies, we also did not include samples that contain potential interfering substances or true samples, however Pastorino *et al* (2020) (27) did include interfering substances and a range of clinical samples and no obvious impact of these sample types were reported on the efficacy of the viral inactivation process.

Due to the contrasting literature for inactivation of SARS-CoV-2 (and that of viruses generally) a case-by-case assessment of different inactivation protocols is essential to prevent laboratory acquired infections. To ensure the highest safety standards (and also taking into account the high viral loads of samples tested), in the operational DCL we employ methods that utilise the inactivation efficacies of the chemical components of lysis buffers found in commercial kits with that of the heat. As a result, the high-throughput RNA extraction platforms are performed on the open bench rather than within a Class 1 microbiological safety cabinet. All laboratories must make the appropriate assessments regarding methods applicable to their unique set of circumstances. The results presented in this study may help laboratories undertake such assessments, especially if they do not have access to high containment facilities to complete in-house inactivation studies.

## ACKNOWLEDGMENTS

The set up and validation of the Defence COVID Laboratory (of which this study was a part) was funded by the UK Department of Health and Social Care (DHSC). The authors thank representatives of Qiagen and ThermoFisher for help in defining suitable RNA extraction protocols.

